# DigiMus: a connectome-informed spiking framework for multi-region mouse neural-behavior modeling

**DOI:** 10.64898/2026.06.09.731075

**Authors:** Yantong Liu, Xinhe Zhang, Xinyu Chen, Chonghe Hao, Wangzi Yao, Jian Zhang, Yue Sun, Tielin Zhang

## Abstract

Computational models are increasingly used to relate mouse brain structure, neural activity and behavior, but most models still learn from task data with limited constraints from biological circuit organization. Here we present DigiMus, a connectome-informed spiking framework for multi-region-capable mouse neural-behavior modeling. DigiMus combines leaky integrate-and-fire spiking dynamics with brain-region-specific motif regularization in a trainable sequence-modeling architecture, allowing directed three-node circuit motifs derived from 38,481 reconstructed neuronal morphologies across approximately 50 brain regions to guide recurrent coupling during learning. We evaluate DigiMus on 18 rule-based cognitive tasks spanning sensorimotor mapping and perceptual decision-making, and on three mouse neural decoding datasets involving auditory discrimination, fixed-interval licking and visual decoding. Across synthetic tasks, DigiMus showed stable performance relative to TCN, LSTM and Transformer baselines, with stronger advantages in more complex decision-making settings. In real neural datasets, single-region instantiations of DigiMus produced small, consistent and dataset-dependent improvements over a structure-free sequence baseline, while retaining motif-prior signatures in trained connectivity. Internal state analyses further linked task-dependent state dynamics to behavioral error patterns. These results suggest that connectome-derived structural priors can shape neural sequence models, and establish DigiMus as a modular, connectome-informed workflow for mouse neural-behavior modeling and hypothesis generation, rather than a complete digital reconstruction.

## Introduction

In recent years, one of the most significant breakthroughs at the biological data level has been the release of diverse brain atlases, including multi-scale data resources such as genomes, connectomes, transcriptomes, and spatial cellular atlases [1,2]. These different types of brain-wide scale biological atlases have provided finer structural and molecular references for brain research, significantly contributing to researchers’ understanding and knowledge of neuronal heterogeneity and brain region functionality [3-5]. In order to better explore the working principle of the mouse brain, a large number of computational models have been proposed one after another, some of which try to align and fuse multi-scale data found during biological experiments, and some of which predict brain activities and brain behaviors from the perspective of computational simulation [6].

However, most of the existing brain activity modeling approaches use generic sequential models such as recurrent neural network (RNN) [7], Transformer [8], or other deep learning architectures for fitting mapping relationships between neural activity, behavior, and external stimuli [9]. While these models have strong end-to-end fitting capabilities, the internal states of the models usually have difficulty in establishing a clear correspondence with neuronal dynamics, synaptic connections, or brain region loop structures, and thus the relationship between their internal representations and the real neural system remains more difficult to interpret [9,10]. In terms of encoding mode, neurons of RNN and Transformer usually rely on static nonlinear activation functions, and do not explicitly describe impulsive neuronal dynamics processes such as membrane potential accumulation, threshold issuance and resetting. In terms of connectivity patterns, the effective connectivity structures of these models are usually learned implicitly by the training process and lack continuous constraints from sparsity, partitioning, and motif topology in the real connectome [11]. As a result, while such models are able to fit the input-output relationship between neural activity and behavior, their internal structure still differs significantly from the neuronal and loop organization in real biological systems, thus limiting their structural traceability and mechanism explanation capabilities. Balancing the processing power of biological signals with the multiscale interpretability of the model’s internal structure is the functional basis of future models for realizing brain simulation and computation. Several studies have attempted to introduce neural loop structures or motif topologies into neural networks to improve the biological plausibility and learning efficiency of models, for example, trying to embed biologically rational network features in the initialization stage [12-15]. Although these methods can bring some learning advantages, if the structural information is mainly used for initialization or weak constraints, the internal connections of the model may still gradually deviate from the original structural settings during the training process. There are also some methods that try to constrain network activity at the level of neural flow shape or population dynamics, such as using Transient trajectory as an additional structural feature to constrain network activity [16,17].

Although these methods make the network reflect some Transient coding effects during the learning process, such constraints usually lack explicit brain region origins and loop topology pointers, thus making it difficult to further support structural-level explanations. Meanwhile, RNNs trained for a particular task often have limited generalization ability and need to be retrained every time they encounter new data or paradigms. Although some models combine continuous learning methods with RNNs [18], they essentially integrate the parameter weights of different models into a single one, and cannot directly solve the problem of the lack of correspondence between the internal structure of the model and the specific loop organization of the brain regions.

Recently, *Drosophila* Cyberfly[19], which is based on the complete connectome of adult *Drosophila*, has successfully driven a virtual body to perform instinctive behaviors in digital space, suggesting that the complete connectome information can provide an important structural basis for animal behavior modeling. However, mouse brain modeling faces a more complex situation, as currently available structural data of the mouse brain are usually derived from cross-scale, heterogeneous, and partially sampled brain atlases rather than complete synapse-level connectomes [20-24]. Neuron-scale structural and connectomic resources of the mouse brain are also being mapped at increasing scale, and are becoming an important source of biological data to support computational modeling of the mouse brain, along with activity and behavioral data [25]. What is special about these connectome data is that the data themselves contain a certain amount of structural knowledge that can be used to constrain the internal connectivity patterns and the organization of information flow within the model, and this structural knowledge is highly relevant to model function, brain region function, and so on [26].

Therefore, an intuitive idea is to use brain-region-specific structural knowledge as biological prior knowledge, and to constrain or incorporate it into neural networks during learning. By allowing structural priors to guide the optimization process, the model may better capture region-dependent circuit organization, rather than relying solely on data-driven parameter fitting. This strategy of incorporating biological structure into model learning differs from traditional purely data-driven models, and the combination of data-driven learning and structural constraints provides a new path for constructing more interpretable neural simulation models. At the same time, the model can also employ spiking neural networks as the underlying computational substrate, introducing dynamics-aware spiking neurons, such as LIF neurons, into the node-level computational process to further improve biological plausibility. Thus, the dual-scale combination of spiking neuron dynamics and connectome-derived motif constraints represents a key direction for constructing an interpretable digital mouse brain modeling framework.

In this study, we first used 38,481 fine neuronal structures obtained after precise tracing with the fMOST technique [27] to analyze projection relationships within and across approximately 50 representative brain regions. We further used a large-scale Golgi-stained mouse-brain morphology resource containing approximately 1.5 million registered neuronal instances. This resource was used as a morphology-based structural reference for estimating regional neuronal distributions and potential local connectivity patterns, rather than as a complete synapse-level whole-brain connectome. Our previous work used this morphology resource to map whole-brain neuronal distributions and identify 12 broad neuronal classes [28]. On this basis, this study proposes DigiMus, a connectome-informed spiking digital mouse brain framework. The available multi-scale brain data are transformed into three modeling operations: LIF-based spiking dynamics, motif regularization and multi-region-capable organization. The core computational unit is the CIS block, namely the connectome-informed spiking block. The CIS block represents LIF-based spiking dynamics at the node level, while connectome-derived motif statistics are used as brain-region-specific motif regularization targets. We further adopt network motif analysis [29,30] to classify possible local connection patterns into 13 connected directed three-node motif classes, and use a unified 13-dimensional vector to represent the network topological features of different brain regions. These brain-region-specific motif vectors are further used to reduce the discrepancy between model-induced local connectivity patterns and connectome-derived statistical targets by applying a motif regularization loss to the low-rank coupling matrix W_r_, forming an interpretable framework with region-specific structural priors. We evaluated DigiMus on an 18-task rule-based cognitive benchmark. For compact visualization, we report six representative tasks covering response, delay, matching and perceptual-decision paradigms, while the remaining 12 tasks were used to assess performance across a broader task set. We further evaluated single-region instantiations of DigiMus on three real mouse neural decoding tasks, including action classification from A1 calcium imaging during auditory discrimination, lick/no-lick classification from M2 extracellular spikes during fixed-interval licking behavior, and drifting-grating direction classification from VISp Neuropixels recordings. DigiMus achieved average accuracies of 72.68%, 78.63% and 70.20% in these three tasks, respectively, and motif-regularized variants outperformed the structure-free Mamba baseline in a dataset-dependent manner. Thus, DigiMus provides a structured computational framework for digital mouse brain modeling by combining LIF-based spiking dynamics, motif regularization and multi-region-capable organization. This framework provides a methodological reference for incorporating connectome-derived structural information into trainable neural sequence models, and a scalable tool for subsequent neuroscience hypothesis generation and computational validation.

## Results

### Multiscale mouse brain data define model inputs

The types of neuroscience experiments vary widely, and the acquired neural signals and structural data are significantly different in spatial scale, temporal scale, and data format depending on the recording devices, experimental means, and subject animals. In this study, we organized AI-ready mouse brain data from resources such as the Brain Data Center and Digital Brain platforms. Neuronal projection structures obtained by fMOST imaging and tracing were normalized into standard SWC-like representations. This format describes key information such as neuronal location, soma size, dendritic branching, axonal projection, synaptic sites, etc. [31,32]. Meanwhile, previous work at the Allen Institute for Brain Research recorded the firing patterns of 1,938 neurons and reconstructed the neuronal morphology of some of them [33]. We aligned different categories of cellular activity by neuronal morphology, and thus fused the two to form a collection of spiking neurons with biological neuronal morphology corresponding to firing activity (**Fig. 1a**).

**Fig. 1.**
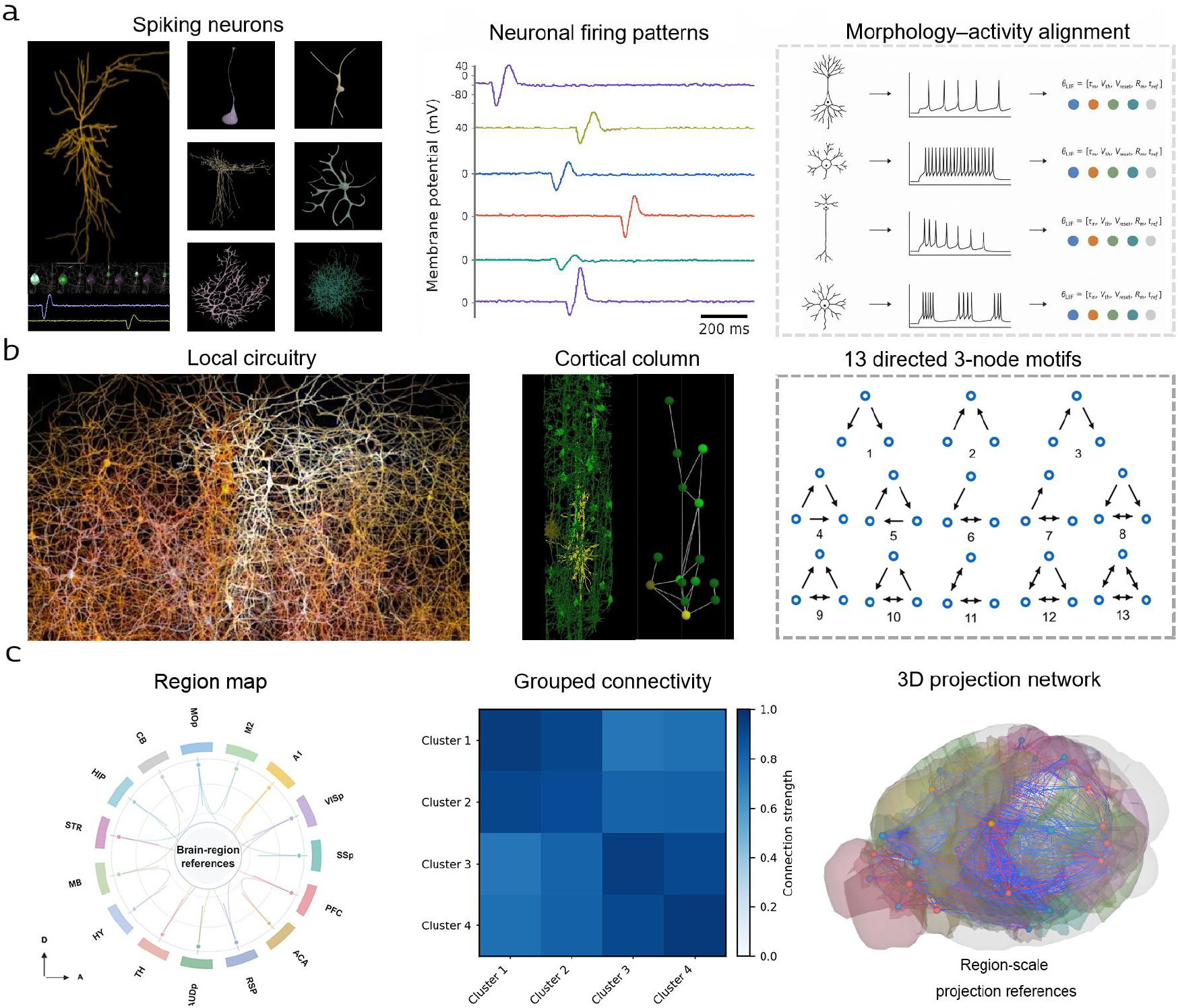
Multiscale neuronal, network and region-level data used to define DigiMus model inputs. **a** Neuron-scale data include spiking neurons, neuronal firing patterns and morphology-activity alignment. fMOST-traced neuronal morphologies were standardized into SWC-like representations, while Allen cell-type electrophysiology data were used to define heterogeneous spiking neuron dynamics. **b** Network-scale data include local circuitry, cortical-column organization and 13 directed three-node motifs. Potential local connections were estimated from spatial relationships between axonal and dendritic structures, and local directed graphs were summarized by motif distributions. **c** Region-scale data include a brain-region reference map, grouped inter-region connectivity and a 3D projection network. Together, the region map, grouped connectivity and projection-network views provide region-level structural references for multi-region-capable organization in DigiMus.

Meanwhile, we summarized the 38,481 neuronal morphologies obtained from the collection and aligned them to standard mouse brain structures. For localized connections within brain regions, the axons and dendrites of these neurons were judged by relative spatial distances to determine whether a potential axon-dendrite contact existed or not. When the shortest axon-dendrite distance was less than a preset threshold and at least one potential axon-dendrite contact was formed, the neuron pair was regarded as having a potential connection; when a certain number of potential contacts was reached, it was further determined that the two neurons were considered to form a directed potential connection. Based on the idea of inferring potential synaptic connections based on neuronal geometric structure, there have been studies on neural geometry and Peters’ rule that provide the methodological basis [34,35]. On this basis, we abstracted the intra-region network as a directed graph structure and used directed three-node motifs as network-scale topological statistics. Without considering the neuron types, directed three-node motifs can be categorized into 13 connected directed three-node motif classes (**Fig. 1b**). These network motif statistics containing fixed categories and statistical frequencies can be used as high-dimensional information statistics in the hidden layer to represent the local topological features of complex networks as 13-dimensional normalized distribution vectors [36,37].

These cells are connected through complex networks to form structures such as nuclei and brain regions, which serve as the data basis to support digital brain simulation. At the brain-region scale, we organized representative brain regions into a region-level reference map and further summarized grouped inter-region connectivity and 3D projection relationships between brain-region groups, resulting in three region-scale representations: a brain-region reference map, grouped connectivity and a 3D projection network (**Fig. 1c**) [38]. These brain-region-scale representations provide structural references for DigiMus’ subsequent region-level organization and potential multi-region routing [39]. On a temporal scale, neural signals characterize the dynamic activity of neurons and neuronal populations in cognitive tasks via high-throughput electrodes, calcium imaging, or other recording modalities.

Traditional AI for Science methods tend to assign pairs of data, such as neural activity, neural structure, neural function, and external behavior, to both model inputs and outputs, or align different modalities through multimodal fusion for intermodal information prediction. Such methods have strong data fitting capabilities, but it is often difficult to establish a clear correspondence between the internal neurons, connectivity structures and state dynamics of the model and the neural data recorded.

In contrast, spiking neural networks introduce neuron representations with spike-based coding and temporal dynamics, and can further incorporate statistical features of connectome-derived local circuits, thus helping to simultaneously balance structural traceability inside the model with data prediction ability outside the model.

### DigiMus integrates spiking dynamics and motif topology

On the basis of the above multi-scale mouse brain data, we designed DigiMus, a digital mouse brain framework based on deep spiking neural networks. First, we mathematically represented neurons with different neurodynamic firing patterns as spiking neuron units containing both spatiotemporal dynamics information and spike-generation properties (**Fig. 2a**). Compared with the static ReLU activation function commonly used in traditional ANN models, LIF-based spiking activation is capable of describing the membrane potential integration, threshold firing and resetting processes, thus introducing a form of pulse encoding closer to neural dynamics in node-level computations (**Fig. 2d**) [40,41].

**Fig. 2.**
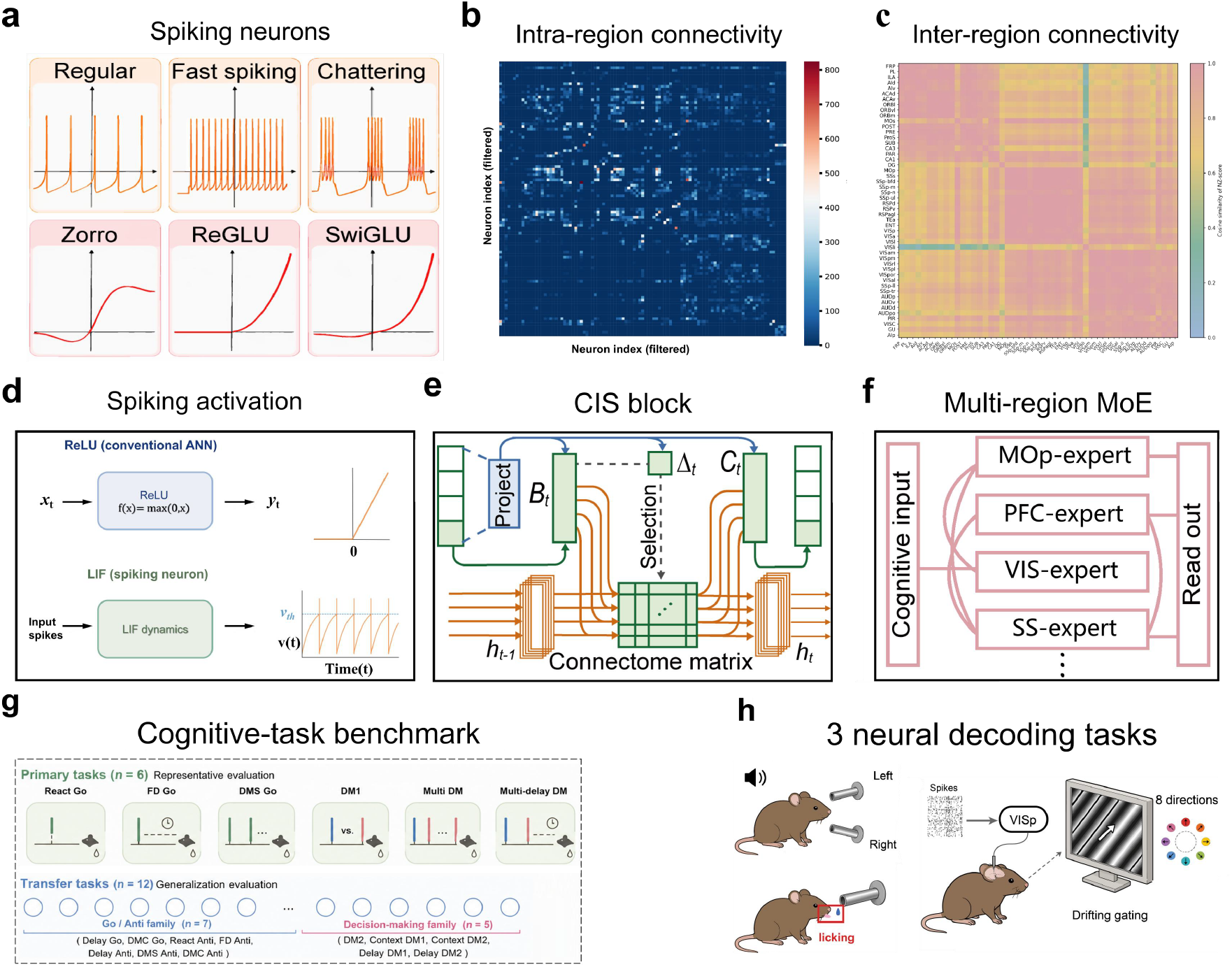
DigiMus framework integrating spiking dynamics, motif regularization and multi-region-capable organization. **a** Spiking neuron units derived from heterogeneous neuronal firing patterns. Different firing modes are mathematically represented as spiking neuron dynamics and used as node-level computational units in DigiMus. **b** Intra-region connectivity matrix showing local connectivity patterns within brain regions. **c** Region-level connectivity reference matrix summarizing sparse inter-region structural relationships. **d** Comparison between conventional ReLU activation and LIF-based spiking activation. LIF dynamics describe membrane potential integration, threshold firing and reset. **e** CIS block architecture. Brain-region-specific motif statistics are used as motif regularization targets to constrain the learnable connectome matrix. **f** Multi-region-capable organization. Region-specific CIS blocks can be instantiated as modular components corresponding to representative brain regions, including MOp, PFC, VIS and SS, and routed or aggregated when multiple brain-region inputs or priors are available. **g** DigiMus is evaluated on 6 primary cognitive tasks and 12 additional tasks. **h** Single-region instantiations of DigiMus are further evaluated on 3 mouse neural-behavior decoding tasks.

At the network structure level, we counted the intra-region and inter-region connectivity maps (**Fig. 2b,c**). Among them, intra-region connectivity patterns were characterized using 13-dimensional motif statistics. Different brain regions have different motif distribution characteristics, so these brain-region-specific motif vectors can be used as motif regularization targets, and the learnable connectome matrix can be continuously regularized by the motif loss function during the network learning process (**Fig. 2e**). This design allows the CIS block to structurally contain statistical features of both spiking neuron dynamics and intra-region connectivity patterns within brain regions.

Meanwhile, sparse connectivity relationships between brain regions are used as structural references for organizing information flow between region-specific modules when multiple brain-region inputs or priors are available. Specifically, different brain regions can correspond to different region-specific CIS blocks, which can be combined through a multi-region-capable routing or aggregation structure when an experiment includes multiple brain-region inputs or priors (**Fig. 2f**) [42]. In this structure, the connectivity relationships between brain regions are not directly equated to the complete biological connectome, but are used as structural references for region-level routing, so that brain-region inputs, outputs and modules can be organized with region-scale structural consistency when multi-region information is available. The framework was organized as a modular workflow in which spiking encoding, motif regularization, region-specific CIS blocks and task-specific readouts can be combined or replaced according to the available neural recordings and structural priors.

These region-specific CIS blocks define the modular components of the DigiMus framework, combining node-level spiking dynamics with local motif-distribution constraints. Specifically, DigiMus was evaluated on six representative cognitive tasks and further evaluated for its broader task coverage in an additional 12 similar tasks (**Fig. 2g**). In addition, single-region instantiations of this framework were evaluated on three real mouse neural-behavioral decoding tasks to assess applicability to real neural recordings and behavioral prediction tasks (**Fig. 2h**). Together, these components define a modeling process from node-level spiking encoding and local motif regularization to region-specific organization, with multi-region routing available when multiple brain-region inputs or priors are provided.

### Brain-region priors support task learning across multiple task types

We selected 18 standardized cognitive computing tasks for evaluating the task learning ability across different cognitive demands of DigiMus under different cognitive demands. These tasks include sensory-motor mapping-like tasks (Go/Anti family) and perceptual-decision-making tasks (DM family) [43]. Among them, there are 10 Go/Anti-family tasks, including reactgo, fdgo, delaygo, dmsgo, dmcgo, reactanti, fdanti, delayanti, dmsanti, and dmcanti, focusing on mapping sensory inputs to behavioral outputs, including both isotropic (Go) and inverse (Anti) responses, and cover different time steps such as immediate response, fixed delay and memory-driven. The The perceptual-decision task family included eight tasks, including dm1, dm2, contextdm1, contextdm2, multidm, delaydm1, delaydm2, and multidelaydm, which involve the comparison and integration of sensory information from multiple sources to accomplish direction selection decisions.

During the information input encoding process, in order to distinguish between different data sources and structural constraint information, we further encoded task-related information such as task category, task text description, relevant brain region information, and cell type as text embedding and spliced it with the neural recording activity data as a unified input representation for the model. This design enables the model to model multi-task information in parallel under the same framework, thus supporting uniform learning and generalization ability assessment across tasks.

In the multiclass cognitive task modeling framework, DigiMus uniformly portrays the mapping relationship between sensory inputs and behavioral outputs. Among them, Go-family tasks mainly generate responses based on a single cue, while DM-family tasks need to compare multiple sources of inputs and select stronger signals, thus reflecting the differences in computational mechanisms across tasks (**Fig. 3a,b**). On this basis, we further constructed a set of representative simulation tasks, including response-type, fixed-delay-type, matching-type, and multimodal decision-type tasks (**Fig. 3c**), to systematically assess the model’s task-learning ability under different cognitive demands.

**Fig. 3.**
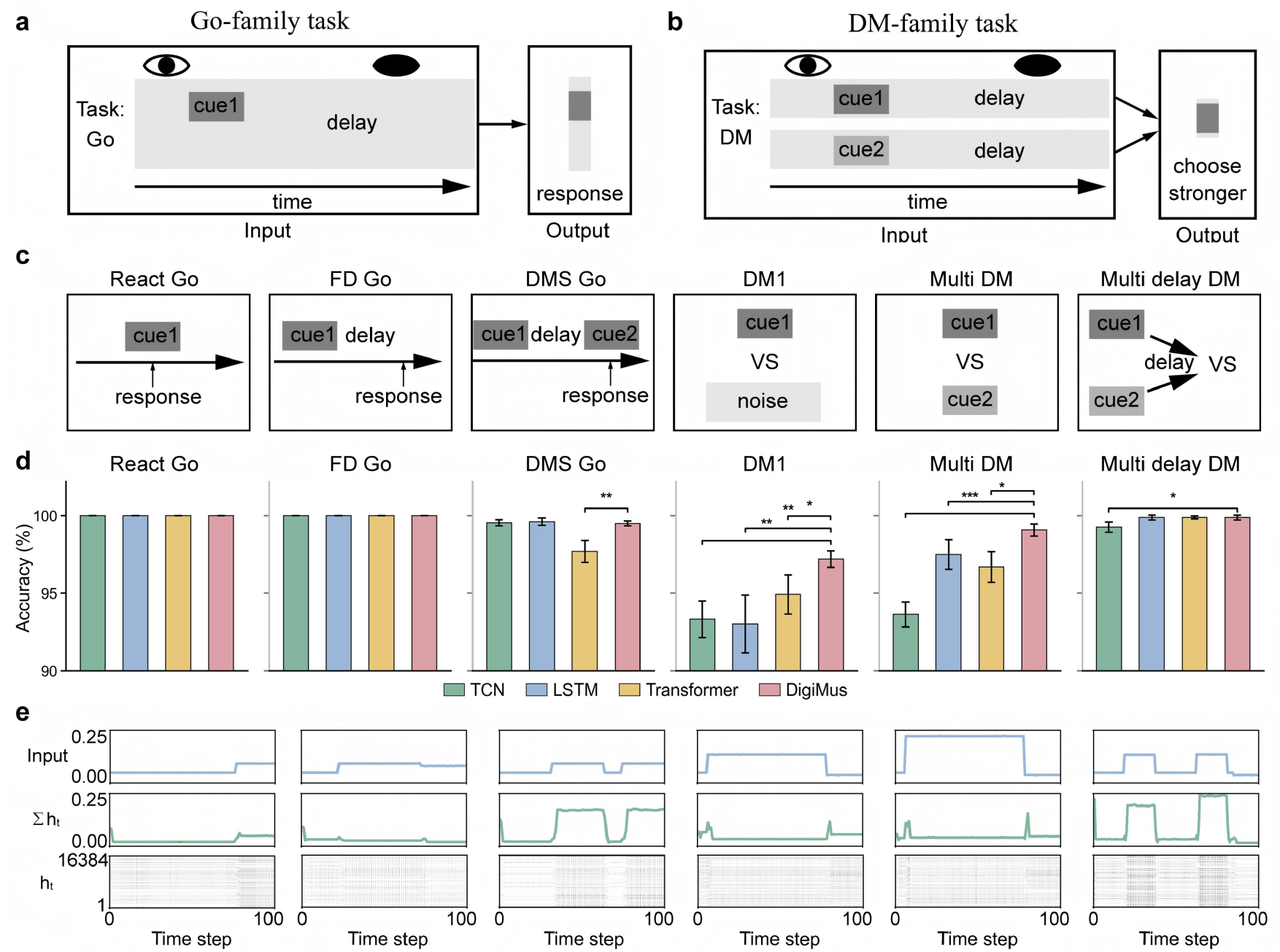
Region-specific constraints support cognitive task learning. **a** Schematic of a Go-family task, in which a sensory cue is mapped to a behavioral response after task-specific timing. **b** Schematic of a DM-family task, in which two sensory inputs are compared and the stronger signal determines the output choice. **c** Representative cognitive tasks used for model evaluation, including React Go, FD Go, DMS Go, DM1, Multi DM and Multi delay DM. **d** Accuracy comparison among TCN, LSTM, Transformer and DigiMus across representative cognitive tasks. Bars show mean accuracy across five random seeds, and error bars indicate SD across seeds. Exploratory model-level comparisons were performed using two-sided Welch’s t-tests comparing each baseline with DigiMus; nominal, uncorrected P values are shown. *P < 0.05, **P < 0.01 and ***P < 0.001. e Internal state dynamics across representative tasks, including task input, population activity I h_t_ and hidden state h_t_ over time.

The experimental results show that DigiMus with motif-regularized brain-region structural constraints exhibits stable and task-dependent behavioral prediction abilities across different types of tasks. In the **Fig. 3** comparison, DigiMus denotes MotifMamba, the motif-regularized Mamba implementation used within the DigiMus framework. Comparisons among individual region priors, including FRP, MOp, CA1 and AUDp, are reported separately when included in the corresponding analysis. DigiMus achieved stable behavioral prediction across the six representative tasks and showed task-dependent mean accuracy gains relative to TCN, LSTM and Transformer baselines, particularly in selected decision-making tasks (**Fig. 3d; Supplementary Table 1**). Analysis of the internal state dynamics of the model further revealed that the input response, group activity, and high-dimensional hidden state evolution of DigiMus exhibit task-related temporal patterns under different task drivers (**Fig. 3e**).

In particular, the input signal reflects the temporal structure of the task stimuli, Σ h_t_ denotes the model population activity over time, and h_t_ denotes the evolution of the high-dimensional hidden state at different time steps. This result suggests that DigiMus is not only capable of accomplishing behavioral output prediction, but also forms dynamic representations that correspond to the temporal structure of the task.

### Task-dependent internal state dynamics

We further analyzed the internal state dynamics of DigiMus in **Fig. 4** across six tasks selected for internal-dynamics analysis to examine whether the model’s internal representations change in response to the computational demands of the task. Unlike comparing only behavioral prediction accuracy, internal state analysis focuses on the hidden-state deviation (*h-dev*) and output-state deviation (*y-dev*) trajectories of the model under different task conditions, which are used to characterize the degree of dispersion and temporal changes of the model state across trial conditions [46,47].

**Fig. 4.**
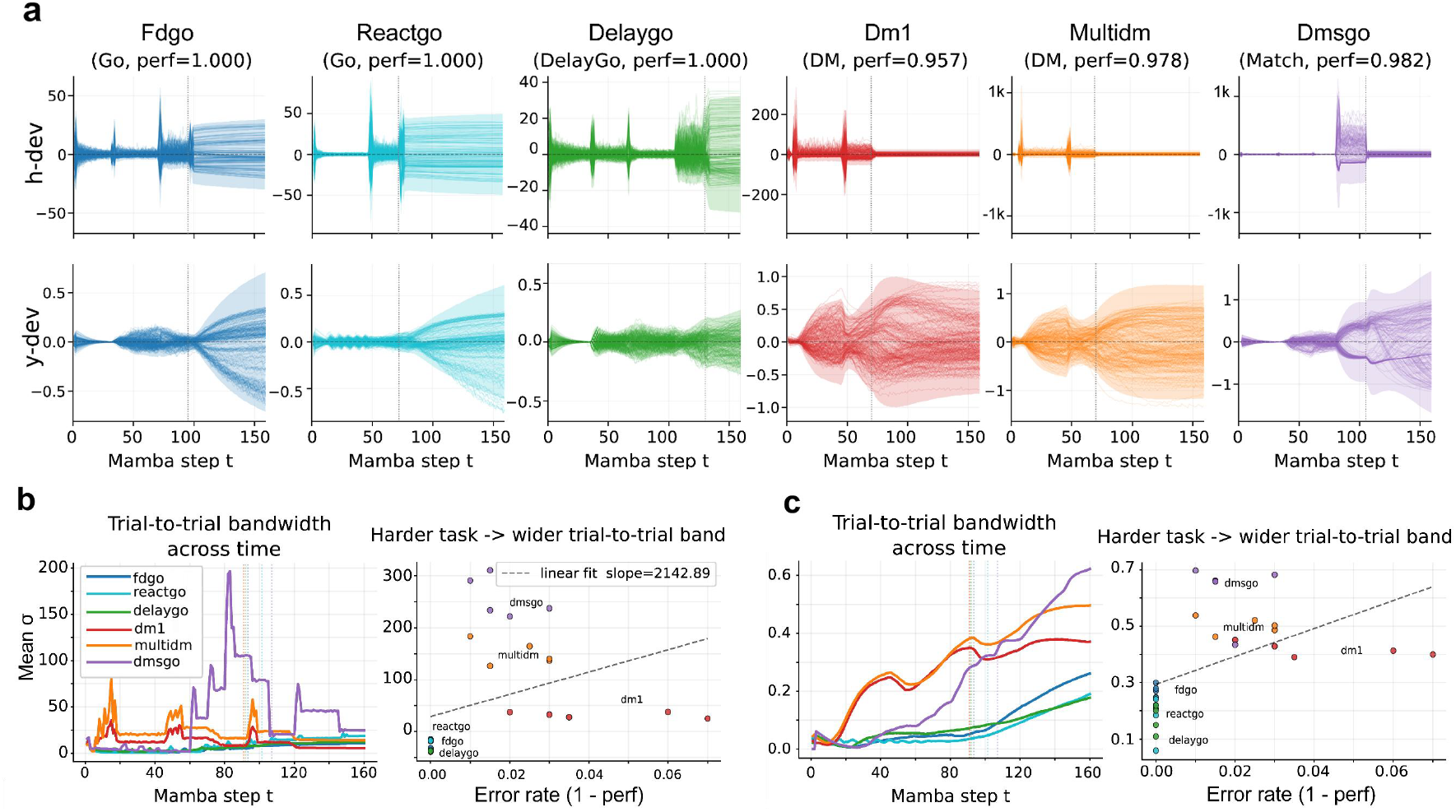
Task-dependent internal state dynamics across representative cognitive tasks. **a** Trial-to-trial trajectories of hidden-state deviation (*h-dev*) and output-state deviation (*y-dev*) across six tasks selected from the same cognitive-task benchmark for internal-dynamics analysis, including FD Go, React Go, Delay Go, DM1, Multi DM and DMS Go. Delay Go included to visualize delay-period state dynamics.. **b** Trial-to-trial bandwidth of *h-dev* across time and its relationship with task error rate. Bandwidth is quantified as the cross-trial standard deviation of state deviation at each time step, and the scatter plot shows task error rate (1™performance) versus peak *h-dev* bandwidth, defined as max σ(t). **c** Trial-to-trial bandwidth of *y-dev* across time and its relationship with task error rate. The scatter plot shows task error rate versus peak *y-dev* bandwidth. Wider trial-to-trial bands indicate larger cross-trial variability in output-state dynamics.

The model achieved high task performance in sensorimotor mapping tasks such as FD Go, React Go, and Delay Go, with corresponding performances shown in the figure as 1.000, 1.000, and 1.000, respectively. Accordingly, the *h-dev* and *y-dev* trial-to-trial trajectories in these tasks were overall more focused, and showed relatively stable temporal state evolution. In contrast, tasks such as DM1, Multi DM, and DMS Go, which involve evidence integration, multi-source input comparison, delayed maintenance, or matching judgment, performed 0.957, 0.978, and 0.982, respectively. *h-dev* or *y-dev* trajectories in these tasks showed a wider cross-trial distribution, which was particularly evident in the task-relevant processing phase (**Fig. 4a**).

Subsequently, we further quantified the trial-to-trial bandwidth in terms of the cross-trial standard deviation of state offsets at each timing step and analyzed its relationship with the task error rate (**Fig. 4b,c**). For the scatter analyses, each task was treated as one effective sample (n = 6), and the peak bandwidth max σ(t) was used as the task-level summary metric. In the *h-dev* analysis, the bandwidth showed task-dependent temporal variation, with some tasks exhibiting higher state dispersion at specific processing stages. The relationship between task error rate and peak *h-dev* bandwidth showed a positive but non-significant trend (slope = 3.0 × 10^3^ ± 3.1 × 10^3^, Pearson *r* = 0.43, *R*^*2*^ = 0.18, *P* = 0.40; Spearman *ρ* = 0.70, *P* = 0.12). In the *y-dev* analysis, output-state bandwidth also varied across time and tasks. The relationship between task error rate and peak *y-dev* bandwidth showed a stronger positive but still non-significant trend (slope = 6.7 ± 4.0, Pearson *r* = 0.64, *R*^*2*^ = 0.41, *P* = 0.17; Spearman *ρ* = 0.70, *P* = 0.12). After Bonferroni correction across the h-dev and y-dev association tests, all corrected P values remained non-significant (corrected P ≥ 0.25; **Supplementary Table 2**).

These results suggest that DigiMus is not only capable of multi-task behavioral prediction, but also forms internal state dynamics related to task timing structure and computational demand. Although the association between task error rate and bandwidth did not reach statistical significance in this six-task analysis, the positive trend suggests that more demanding tasks may induce broader trial-to-trial state dispersion. The higher *h-dev* or *y-dev* bandwidth may reflect greater state uncertainty during evidence integration, delay maintenance and matching judgment.

### Region-specific decoding across mouse neural datasets

To examine whether DigiMus can be applied beyond synthetic cognitive tasks, we evaluated the motif-regularized sequence modeling framework on three mouse neural decoding datasets spanning A1 calcium imaging, M2 extracellular spike recordings and VISp Neuropixels recordings. In these real-data experiments, neural activity from task-relevant single-region recordings was mapped to behavioral or stimulus labels, while the same motif-regularized sequence modeling framework was preserved. These experiments were designed to test whether connectome-derived motif constraints could improve neural decoding performance across different recording modalities and task structures.

The first dataset involved an auditory frequency discrimination task in head-fixed mice. Neural activity was recorded from the primary auditory cortex (A1) using two-photon calcium imaging while mice reported low- or high-frequency pure-tone stimuli through left or right licking responses. For each trial, the model used the post-stimulus neural activity tensor within a 1 s window as input and predicted the behavioral action label from post-stimulus neural activity. This task was used to evaluate whether DigiMus could model the transformation from sensory evidence to behavioral choice in calcium-imaging recordings (**Fig. 5a**)[48].

**Fig. 5.**
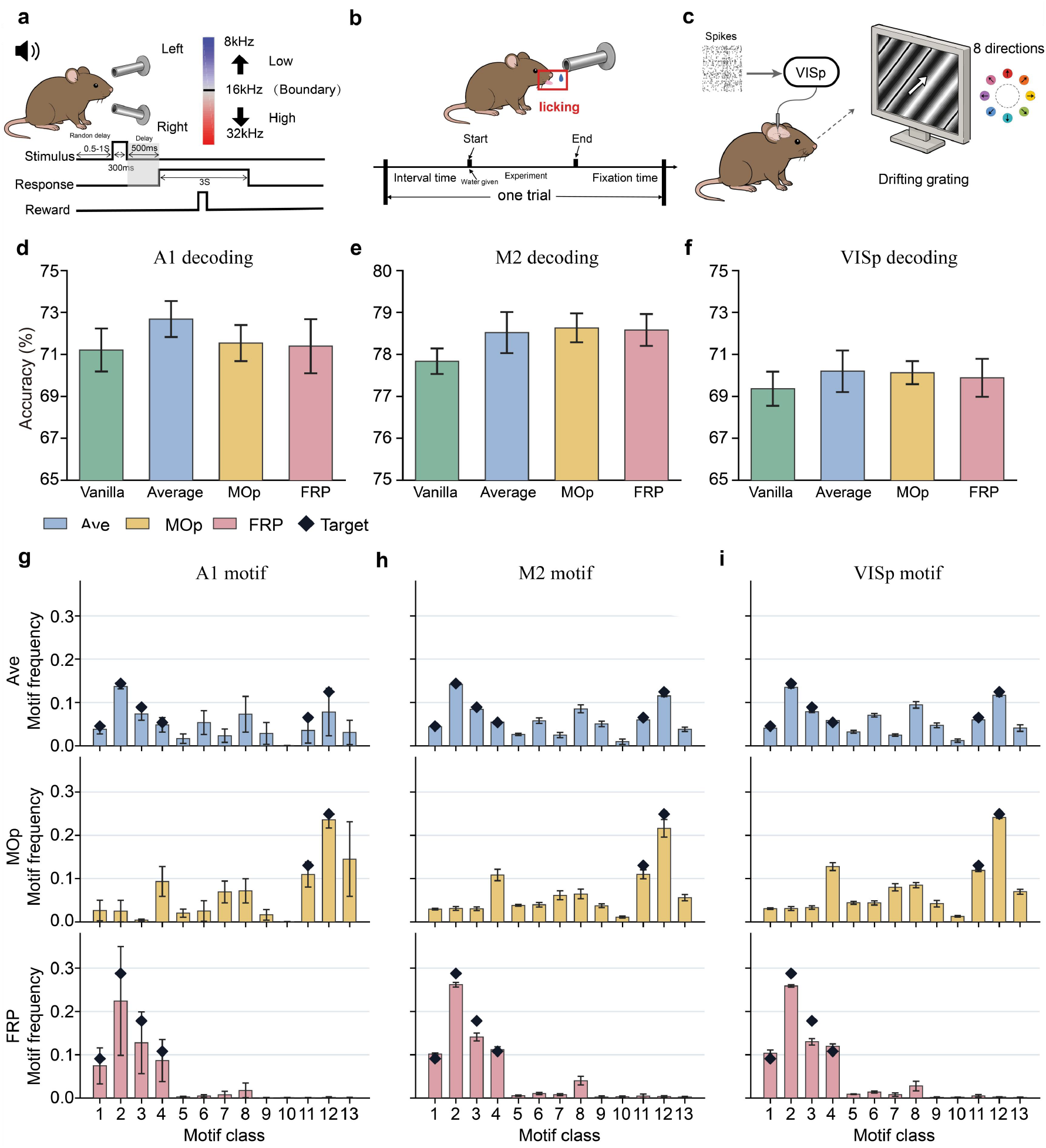
DigiMus generalizes to mouse neural-behavior decoding datasets. **a** A1 calcium-imaging action classification task. Mice performed an auditory discrimination task and reported their choices through left or right licking responses; the model used post-stimulus A1 calcium activity to predict the behavioral action label. **b** M2 fixed-interval licking task using extracellular spike recordings from the M2 subset. Trial-aligned spike sequences were used to predict licking events across time windows. **c** Drifting-grating direction decoding task based on VISp Neuropixels recordings from awake mice during passive viewing. **d** Decoding performance on the A1 calcium action-classification dataset. **e** Decoding performance on the fixed-interval licking dataset. **f** Decoding performance on the drifting-grating direction decoding dataset. **g-i** Motif-frequency distributions of trained model-induced recurrent connectivity under Average/Ave, MOp and FRP motif constraints across the three decoding datasets, including the A1 calcium action-classification dataset (**g**), M2 fixed-interval licking dataset (**h**) and VISp drifting-grating direction decoding dataset (**i**). Colored bars indicate model-induced motif-frequency distributions after training, and black diamonds indicate the corresponding target motif-prior profiles.

The second dataset involved fixed-interval licking behavior in water-restricted, head-fixed mice. Although the source dataset contains extracellular spike recordings from multiple brain regions, including M2, VLS and SNR, the present analysis was restricted to a single-day M2 recording subset. DigiMus used trial-aligned M2 spike sequences as input and predicted whether licking occurred within each time window. This experiment was therefore formulated as a single-day M2 lick/no-lick classification task and was used to evaluate whether motif-regularized sequence modeling could improve decoding of temporally structured licking behavior from motor cortical spiking activity(**Fig. 5b**)[49].

The third dataset was based on the Allen Brain Observatory Visual Coding Neuropixels resource. We used extracellular spiking activity from the primary visual area (VISp) recorded while awake mice passively viewed drifting gratings presented in eight motion directions. The model used population spike trains or firing-rate sequences as input and predicted the stimulus direction for each trial. This task was used to evaluate whether DigiMus could decode visually evoked cortical activity under a multiclass stimulus classification setting (**Fig. 5c**)[50].

In the A1 calcium action-classification dataset, motif-regularized DigiMus variants showed modest but consistent improvements over the structure-free Mamba baseline. Vanilla Mamba achieved an average accuracy of 71.21%, whereas Ave, MOp and FRP motif-frequency constraints reached 72.68%, 71.54% and 71.39%, respectively. Among these variants, Ave obtained the highest mean accuracy, improving over vanilla Mamba by approximately 1.47 percentage points (**Fig. 5d**). These results suggest that motif regularization can modestly improve decoding performance for calcium-imaging-based action classification, although the magnitude of improvement remains moderate.

In the single-day M2 fixed-interval licking dataset, all three motif-regularized variants outperformed vanilla Mamba. The baseline achieved an average accuracy of 77.84%, whereas Ave, MOP and FRP reached 78.52%, 78.63% and 78.58%, respectively (**Fig. 5e**). MOp produced the highest average performance, while FRP and Ave also exceeded the baseline. However, this task showed substantial class imbalance, with no-lick windows accounting for approximately 74.58% of samples. Accuracy alone could therefore overestimate decoding performance. Additional diagnostic metrics showed a balanced accuracy of approximately 0.67–0.68 and a lick recall of approximately 0.46–0.47. These results indicate that the model did not collapse to majority-class prediction, but lick-event detection remained challenging. Thus, in the single-day M2 licking task, motif regularization should be interpreted as providing a stable, moderate improvement in within-day lick/no-lick decoding rather than fully resolving the lick/no-lick imbalance.

In the VISp drifting-grating direction decoding dataset, the improvement from motif regularization was smaller than that observed in the other two datasets. Across five random seeds, vanilla Mamba achieved an average accuracy of 69.36%, whereas Ave, MOp and FRP reached 70.20%, 70.13% and 69.88%, respectively (**Fig. 5f**). Ave obtained the highest mean accuracy, improving over vanilla Mamba by approximately 0.83 percentage points. MOp and FRP also exceeded the baseline, but their gains were smaller. This suggests that the benefit of motif regularization depends on recording modality, task structure and dataset heterogeneity. In particular, the visual decoding task involved more stimulus classes and stronger session-level variability, which may reduce the effect size of a fixed motif-frequency constraint.

Across the three real neural decoding datasets, motif-regularized variants consistently improved over the structure-free Mamba baseline. No single motif prior dominated across all neural decoding datasets, indicating that the effective structural constraint depends on recording modality, task structure and readout target. Ave achieved the highest accuracy in the A1 calcium action-classification task and the VISp drifting-grating direction decoding task, whereas MOp achieved the highest accuracy in the single-day M2 fixed-interval licking task. These results indicate that connectome-derived motif regularization can be incorporated into sequence models of mouse neural activity without degrading decoding performance, and may provide small but consistent improvements in selected neural decoding settings. Importantly, the task-dependent variation across Ave, MOp and FRP suggests that motif constraints are not universally interchangeable and should be evaluated with respect to the neural modality and task readout under study.

### Model-to-data alignment in virtual mouse simulation

Beyond decoding behavioral variables from recorded neural activity, we further examined whether the motif-regularized DigiMus variants retained their prescribed structural priors after training on real mouse neural-behavioral datasets. In this analysis, the model-induced recurrent connectivity was summarized as directed three-node motif-frequency distributions and compared with the corresponding motif-prior targets used during training. This analysis was designed to evaluate whether motif regularization remained visible in the trained model structure, rather than only producing changes in decoding accuracy.

First, we compared the motif distribution of the trained model-induced intra-region connectivity with the target motif-frequency profiles defined by Ave, MOp and FRP priors. Across the three neural decoding datasets, the induced motif distributions generally showed prior-dependent convergence patterns, with Ave-, MOp- and FRP-constrained models tending to preserve different motif-frequency signatures after task training (**Fig. 5g-i; Supplementary Table 3**). Thus, **Fig. 5g-i** should be interpreted as a structural-retention analysis of motif regularization, rather than as a full biological validation of mouse brain connectivity.

Second, we examined whether these motif-constrained variants maintained neural-behavioral decoding performance across different recording modalities. The A1 calcium-imaging, M2 extracellular-spiking and VISp Neuropixels datasets differed in signal type, brain region and prediction target, but motif-regularized variants achieved small and dataset-dependent improvements over the structure-free Mamba baseline (**Fig. 5d-f**). These results suggest that motif constraints can be incorporated into neural sequence models without degrading decoding performance, although the magnitude of improvement remains moderate and task-dependent.

Third, we assessed whether the best-performing motif prior was consistent across datasets. The best-performing prior differed across neural decoding settings, with Ave showing stronger performance in the A1 and VISp datasets, whereas MOp showed the highest mean accuracy in the M2 licking dataset. This pattern suggests that motif priors are not universally interchangeable and that their effects may depend on recording modality, task structure, regional origin and behavioral readout.

Together, these analyses provide a conservative model-to-data consistency view of DigiMus. Rather than claiming that the trained model structure or activity fully reproduces recorded biological neural dynamics, the current **Fig. 5** analysis shows that motif-regularized DigiMus can retain prescribed structural priors while maintaining or modestly improving neural-behavioral decoding performance. This result supports the use of connectome-derived motif regularization as a structural constraint for mouse neural sequence modeling and provides a basis for future analyses that directly compare model activity trajectories with recorded neural population dynamics.

## Connectome-informed visualization layer for model inspection

Beyond model training and neural-behavioral decoding, we developed a connectome-informed visualization and simulation interface to make the internal organization of DigiMus inspectable in anatomical, structural and behavioral coordinates (**Fig. 6**). This interface was not intended as a conventional graphical display module, but as a structure-function inspection layer linking the main components of the DigiMus framework: brain-region priors, region-specific model configuration, model-generated activity summaries and behavior-related outputs. By placing these components within a unified visualization workflow, DigiMus provides an explicit route for examining how connectome-derived structural priors are assigned, how model states are organized in brain-region space and how simulated outputs are related to task-level behavior.

**Fig. 6.**
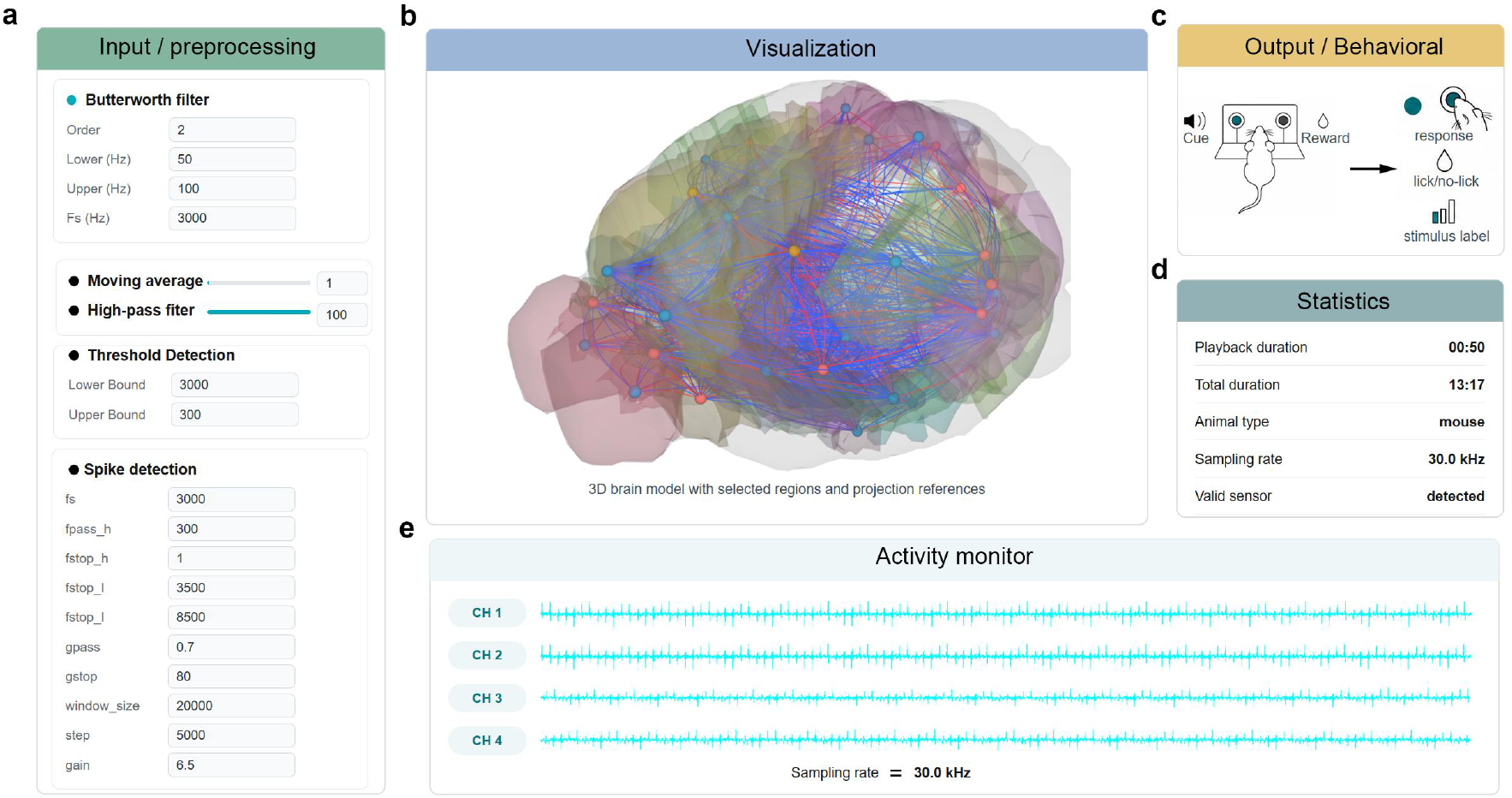
Connectome-informed visualization layer for DigiMus inspection. **a** Input and preprocessing panel showing representative signal-processing and spike-detection controls used in the visualization workflow. **b** Three-dimensional visualization of registered mouse brain regions with selected projection references. **c** Behavioral-output panel linking task cues, reward events and model-generated responses. **d** Diagnostic summary panel displaying representative simulation and recording metadata. **e** Activity-monitor panel showing multi-channel neural activity traces used for visual inspection. The interface was designed as a diagnostic layer for DigiMus, allowing users to inspect preprocessing settings, brain-region organization, projection references, simulated activity and behavior-related outputs within a unified graphical environment.

The interface was organized into five functional views. The input and preprocessing view provides access to signal-processing, smoothing, threshold-detection and spike-detection settings used before model inspection (**Fig. 6a**). The brain-region visualization view displays a registered three-dimensional mouse-brain representation with selected region and projection references, allowing anatomical organization and inter-regional structural relationships to be inspected in a common visual space (**Fig. 6b**). The behavioral-output view links task cues, reward-related events, response outputs and stimulus labels, providing a compact representation of how model-generated outputs can be related to task-level behavior (**Fig. 6c**). The statistics view summarizes essential simulation and recording metadata, including playback duration, animal type, sampling rate and sensor status (**Fig. 6d**). The activity-monitor view displays multi-channel activity traces, providing a temporal entry point for inspecting activity patterns during model visualization and simulation monitoring (**Fig. 6e**). The original software interface is shown in **Supplementary Fig. 1**.

We further used the interface to generate structure-function visualization outputs for mouse-brain representations (**Fig. 7**). These outputs include registered atlas views, rotational views of brain-region structures, sectional activity projections, whole-brain projection references and spatial activity-map visualizations. Together, these panels illustrate how anatomical references, projection networks and DigiMus-associated activity summaries can be inspected within the same visual framework. These visualizations should not be interpreted as evidence that DigiMus reproduces the full biological mouse brain. Rather, they demonstrate that the framework can organize structural priors, model states and behavioral readouts into an inspectable simulation-facing environment. Additional simulated mouse-brain visualization assets and registered atlas views are shown in Supplementary **Fig. 2** and **Fig.3**.

**Fig. 7.**
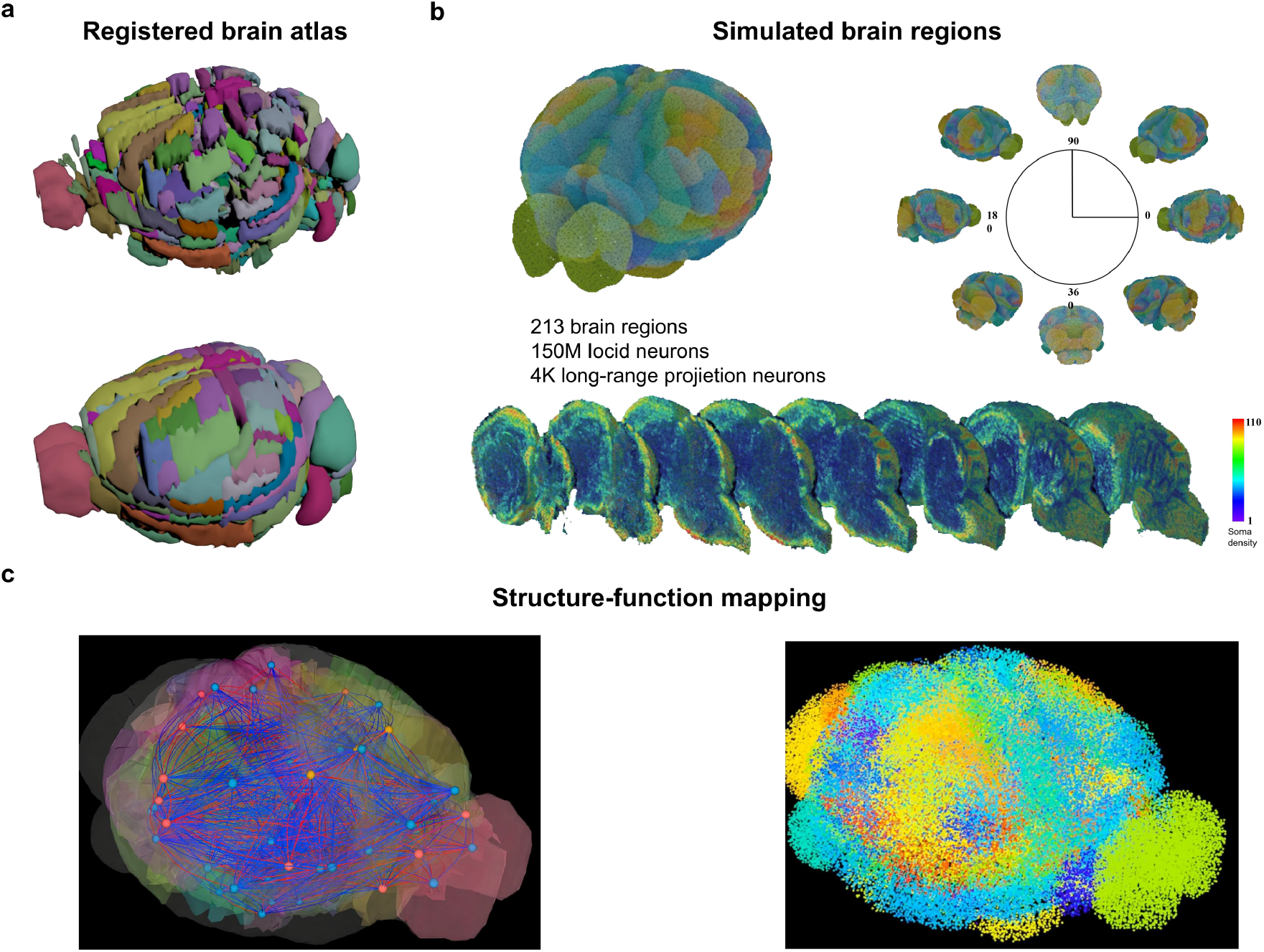
Structure-function visualization of simulated mouse-brain representations. **a** Registered mouse-brain atlas views showing region-level parcellation used for anatomical visualization. **b** Simulated mouse-brain region representation with rotational views and sectional activity projections. **c** Structure-function visualization view displaying region-level projection references together with simulated activity maps. These visualizations illustrate how DigiMus-associated anatomical references, projection networks, motif-related structural priors and model-generated activity summaries can be displayed for figure-level inspection and simulation monitoring.

This visualization layer extends DigiMus from a trainable sequence model into an inspectable digital mouse-brain modeling environment. In contrast to purely data-driven neural decoders, whose internal representations are usually examined only through post hoc analysis, DigiMus allows model configuration, anatomical references, state dynamics and behavioral outputs to be inspected within the same connectome-informed workflow. This design supports model diagnosis, figure-level visualization and future closed-loop validation with richer neural recordings and perturbation data.

## Discussion

DigiMus was developed as a connectome-informed spiking framework for linking mouse brain structure, neural dynamics and behavior within a unified computational model. Rather than treating neural-behavior modeling as a purely input-output mapping problem, DigiMus incorporates two levels of biological prior knowledge: node-level spiking dynamics and region-specific motif topology. This design allows the model to retain explicit computational elements that can be related to neuronal firing, local circuit organization and potential inter-region structural routing. Across synthetic cognitive tasks and mouse neural-behavior decoding datasets, the present results suggest that such biological priors can be incorporated into sequence models without sacrificing task performance, while providing a more interpretable internal organization than structure-free deep learning models [6,9]. In this sense, DigiMus should be viewed not as a complete digital replica of the mouse brain, but as a biologically constrained modeling framework for testing how structural priors may shape neural computation.

A central feature of DigiMus is the use of motif statistics as a compact representation of brain-region-specific circuit organization. Network motifs have long been considered recurring local building blocks of complex biological networks, including neural systems [11,14,37,51,52]. Compared with simple degree, hub or small-world measures, directed three-node motifs provide a higher-order description of local connectivity patterns and may capture circuit properties that are relevant to information integration, recurrent amplification and state stabilization. In DigiMus, motif distributions are not used merely as descriptive statistics, but as regularization targets that constrain the learnable connectome matrix during training. This provides a practical route for embedding connectome-derived structural knowledge into trainable neural models. However, motif regularization remains a coarse approximation of biological circuitry. It does not yet capture synaptic weights, cell-type-specific connectivity, dendritic integration, neuromodulation or activity-dependent plasticity. Future versions of DigiMus should therefore integrate motif priors with transcriptomic, electrophysiological and synapse-level resources as these data become more complete [53,54].

The performance of DigiMus across 18 cognitive tasks and three neural-behavior decoding datasets suggests that biological structure and computational flexibility can be combined within a single modeling framework. The synthetic tasks show that region-specific structural priors can support diverse cognitive computations, whereas the real mouse datasets indicate that motif-regularized variants can achieve modest but reproducible gains in selected decoding settings. Importantly, these improvements are dataset-dependent rather than universal. This is a strength of the current interpretation: the results do not imply that one fixed motif prior is optimal for all neural modalities or behavioral tasks. Instead, they suggest that different structural priors may interact with recording modality, task complexity and behavioral readout. The internal-state analyses further indicate that trial-to-trial state dispersion varies across task classes, consistent with the broader view that neural computation is organized through task-dependent population dynamics rather than static feature extraction [10,55–57].

Several limitations define the current scope of DigiMus. First, the neuron types used in the model are still simplified representations derived from morphology and electrophysiology. Real neuronal identity is shaped by morphology, firing pattern, transcriptomic state, projection target and developmental history, and future digital brain models will need to incorporate these axes jointly [53,58]. Second, the structural connectome used here is inferred from available morphology- and atlas-derived resources rather than a complete synapse-level mouse brain connectome. Potential contact-based connectivity and motif statistics provide useful structural priors, but they should not be interpreted as direct measurements of all biological synapses. Third, the present model-to-data alignment analyses remain preliminary and should be strengthened with quantitative comparisons of motif similarity, neural activity trajectories and behavioral output consistency across trials, sessions and animals. Finally, prospective experimental validation will be needed to test whether model-derived region or motif predictions can guide new neural recordings, perturbation experiments or task designs.

Together, these findings support DigiMus as a step toward biologically constrained digital mouse brain modeling. Its main contribution is not simply higher decoding accuracy, but the construction of a modeling framework in which spiking dynamics, connectome-derived motif priors and multi-region-capable organization are jointly represented. This framework provides a white-box alternative to purely data-driven neural-behavior models and offers a platform for comparing model activity with biological structure and behavior in a measurable way [6,9]. As richer mouse brain atlases, neural recordings and connectomic datasets become available, DigiMus can be extended from the current motif-regularized framework toward more detailed, testable and experimentally grounded digital brain simulations.

## Methods

### Data resources and prior digital-brain platforms

DigiMus was developed as a computational modeling framework using synthetic cognitive tasks, retrospective mouse neural-behavioral recordings, and publicly available mouse brain morphology, electrophysiology and connectome-related resources. No new neural recording, behavioral training, surgery or prospective animal experiment was conducted specifically for this study. Instead, the present work focused on reorganizing existing brain data resources into a trainable connectome-informed spiking framework.

The data organization and structural-prior design of DigiMus were informed by public brain data and digital-brain resources, including the Brain Science Data and Computing Center of the Center for Excellence in Brain Science and Intelligence Technology, Chinese Academy of Sciences, and the Digital Brain / Mouse Brain Atlas platforms [59-61].

These resources provide important references for brain data management, computational analysis, atlas-based visualization, neuronal reconstruction and region-level structural organization. In DigiMus, these resources were not used to claim a complete biological replica of the mouse brain, but to support the construction of a structurally constrained computational framework in which brain-region priors, neuronal dynamics and neural-behavioral decoding can be jointly represented.

DigiMus is also positioned in relation to prior digital-brain and digital-twin-brain modeling efforts. Earlier digital-brain platforms emphasized the integration of brain databases, computational brain phantoms, knowledge bases and intelligent research assistants for brain research [62]. More recent digital twin brain frameworks have further proposed the use of brain structure, bottom-level generative models and application-oriented simulations to bridge biological intelligence and artificial intelligence [63]. In addition, large-scale brain models constrained by multimodal neuroimaging data have shown that computational simulations can reproduce experimentally observed whole-brain response patterns, such as steady-state visual evoked potentials, and can be used to investigate the dynamic mechanisms underlying brain activity [64,65].

Compared with these previous digital-brain and human neuroimaging-based digital twin brain studies, DigiMus focuses specifically on mouse neural-behavioral modeling. Its main methodological distinction is the integration of LIF-based spiking dynamics, connectome-derived directed three-node motif regularization and multi-region-capable model organization. Thus, DigiMus should be interpreted as a connectome-informed computational framework for testing how mouse brain structural priors may shape neural sequence modeling, rather than as a complete digital reconstruction of the biological mouse brain.

### Cognitive task datasets

The synthetic cognitive-task datasets were generated from rule-based task paradigms commonly used in computational cognitive neuroscience. The task set contained 18 simulated cognitive tasks covering sensory-guided response, delayed response, rule-dependent mapping and perceptual decision-making processes. These tasks were designed to provide controlled benchmarks for evaluating whether DigiMus could learn diverse input-output mappings while maintaining biologically constrained internal dynamics.

The 18 tasks were grouped into two broad families according to their dominant computational demands. The first family comprised sensorimotor mapping tasks, including Go and Anti-Go variants. In these tasks, the model was required to transform sensory input into an appropriate behavioral response, either in the same direction as the stimulus or in the opposite direction. This family further included different temporal structures, such as immediate reaction, fixed-delay and memory-guided response tasks. The second family comprised perceptual decision-making tasks, in which the model compared or integrated sensory evidence across one or more input modalities before generating a choice. Some tasks additionally included delay periods, requiring the model to maintain task-relevant information before response generation. The general task structure followed established multi-task cognitive modeling paradigms [43,55].

All tasks used a unified trial-based input-output format. Each input sequence contained three components: a fixation input indicating whether the model should maintain a waiting state or enter the response period; sensory stimulus inputs consisting of two sensory modalities, each represented by a population of ring-tuned units encoding continuous stimulus variables such as direction; and a task-rule input specifying the task identity for the current trial. The output sequence contained a fixation output and a continuous response-direction output, which together defined the expected behavioral response.

Each trial consisted of temporally structured epochs, including a fixation period, stimulus period, optional delay period and response period. For tasks without a memory component, the model could generate the behavioral response after stimulus presentation. For delay or decision-making tasks, the model was required to maintain or integrate task-relevant information across time before producing the correct output during the response epoch. Training targets were generated according to the rule of each task, and task performance was evaluated by comparing the model output with the corresponding target response during the response period.

### Mouse Neural-Behavioral Datasets

DigiMus was evaluated on three mouse neural-behavioral decoding datasets that differed in recording modality, brain region and behavioral or stimulus paradigm. These datasets were used to examine whether the connectome-informed spiking framework could be applied beyond synthetic cognitive tasks to neural-to-behavior or neural-to-stimulus decoding under heterogeneous experimental conditions.

The first dataset was an auditory two-alternative forced-choice dataset with simultaneous two-photon calcium imaging from primary auditory cortex (A1) and behavioral recordings [48]. In the original task, mice categorized auditory stimuli as low- or high-frequency tones and reported their choices by licking the left or right water port. For the experiments reported here, trial-aligned A1 calcium activity was used as the neural input, and only the behavioral action label was used as the supervised target. Frequency and reward labels were not used as prediction targets in the present analysis. This dataset was therefore used to evaluate action classification from A1 calcium-imaging recordings.

The second dataset contained aligned neural spiking activity and licking behavior recorded from thirsty mice [49]. Although the source dataset includes recordings from M2, VLS and SNR, the experiments reported here used only the M2 subset. Trial-aligned M2 spike sequences were used as model inputs, and lick/no-lick labels were used as decoding targets. This dataset was used to evaluate whether DigiMus could predict temporally structured licking behavior from M2 population spiking activity. Train, validation and test partitions were generated within the M2 subset according to the preprocessing protocol used for the reported experiments.

The third dataset was derived from the Allen Brain Observatory Visual Coding Neuropixels resource [50]. We used spike-train or firing-rate sequences from selected visual cortical recordings during drifting-grating stimulation. The supervised target was defined as the stimulus direction category. When the analysis was restricted to primary visual cortex, units were selected from VISp before model training. This dataset was used to evaluate stimulus decoding from visually evoked cortical population activity under a multiclass classification setting.

For all three datasets, neural activity was converted into trial-aligned temporal sequences before model training. Dataset-specific preprocessing included temporal window selection, binning or resampling of neural activity, label extraction, normalization and train-validation-test splitting. For each dataset, all compared variants used the same preprocessing, train-validation-test split, input sequence format and supervised target; only the motif-prior constraint differed between motif-regularized variants and the structure-free Mamba baseline. Model performance was evaluated using the metrics reported for each dataset. Accuracy was used for balanced multiclass decoding tasks, whereas imbalance-aware metrics, including balanced accuracy and event recall, were additionally reported for licking-behavior decoding when class imbalance was present.

### Model architecture and training

We next describe the DigiMus model components used for these decoding and cognitive-task experiments. The framework combines LIF-based spiking encoding, connectome-informed CIS blocks, motif regularization, multi-region-capable organization and task-specific readout layers within a unified trainable sequence-modeling pipeline.

### LIF spiking encoding

For any input time series x_t_∈R^d^, we first map it to the input current I_t_, of the spiking neuron. The membrane potential of each spiking neuron is updated according to the LIF-based spiking dynamics:

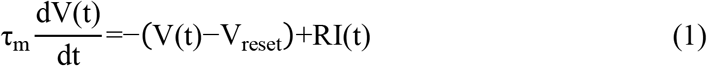

where τ_m_ is the membrane time constant, V(t) is the membrane potential at moment t, Vreset is the reset potential, *R* is the membrane resistance, and I(t) is the external input current. The spiking neuron generates a pulse when the membrane potential exceeds the threshold:

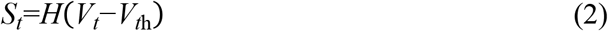

where *H*(⋅) denotes the Heaviside step function. When *S*_*t*_=1, the membrane potential was reset to V_reset_.

Because the threshold function is non-differentiable, we used a surrogate gradient for back-propagation:

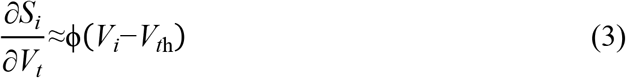

where ϕ(⋅) denotes the surrogate derivative. In the current implementation, ϕ(⋅) was set as the derivative of a sigmoid function. After LIF encoding, the continuous input sequence was converted into a spike sequence:

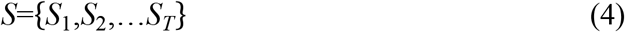

### CIS block architecture

After LIF encoding, the spike sequence was passed into a connectome-informed spiking (CIS) block for temporal modeling. The CIS block used a Mamba-style state-space temporal backbone and introduced a low-rank recurrent coupling term to support cross-channel state interaction. Let the input spike sequence be *u*_*t*_ and the hidden state be h_*t*_∈*R*^*n*^. The standard discrete state update was written as:

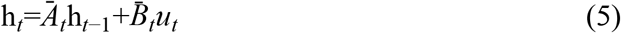

where 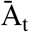and 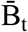 denote input-dependent state transition and input projection matrices.

To enhance information exchange across hidden dimensions, we added a recurrent coupling matrix 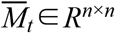:

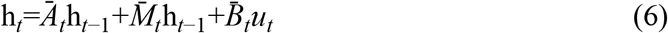

To control parameter size and computational cost, 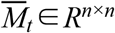 was parameterized in low-rank form:

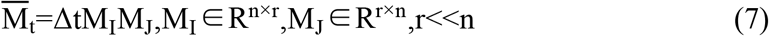

This low-rank coupling can be viewed as a cross-channel temporal mixing mechanism that preserves the efficiency of the state-space backbone while adding structured recurrent interaction.

### Motif regularization

To incorporate connectome-derived structural information, the recurrent coupling matrix was treated as an induced directed weighted graph. From this graph, we computed the distribution of directed three-node motifs:

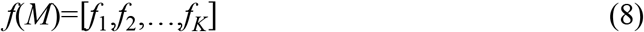

Here, *K*=13 denotes the 13 connected directed three-node motif classes used in this study, and f(M) describes the motif-count distribution induced by the model-internal recurrent structure.

For each brain region, a target motif vector f^*^ was estimated from the corresponding connectome-derived regional topology statistics. The motif loss was defined as the discrepancy between the model-induced motif distribution and the biological target motif distribution:

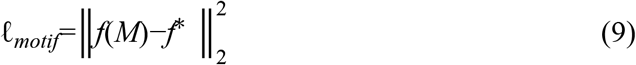

This constraint was designed to preserve local connection-pattern information while allowing the model to learn task-relevant temporal dynamics.

### Multi-region organization

DigiMus can be instantiated with multiple region-specific CIS blocks when an experiment includes inputs or priors from more than one brain region. When only one recorded brain region was available, the aggregation operator reduced to the identity mapping and the model was evaluated as a single-region DigiMus instantiation with the corresponding motif prior. For a modeled region r, the corresponding regional block E_r_ receives the region-specific input x_r_ and produces a latent temporal representation:

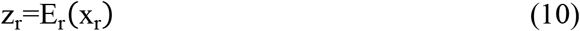

The structural prior of each regional block is defined by the motif statistics of the corresponding brain region, allowing different blocks to encode region-dependent local topology.

When multiple region-specific CIS blocks were used, their outputs were integrated by the aggregation rule implemented for the corresponding experiment:

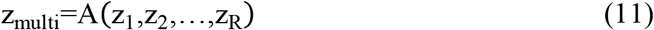

Where R denotes the number of modeled brain regions and A(·) denotes the code-level aggregation operation.

The integrated representation was then passed to a task-specific readout head for behavioral, stimulus or outcome prediction.

### Training objective

When motif regularization was used, the training objective was written as:

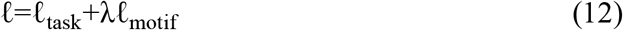

where ℓ_*task*_ denotes the supervised loss for the corresponding behavioral or decoding task, ℓ_motif_ denotes the motif-constrained loss and *λ* controls the strength of the structural constraint.

For synthetic cognitive tasks, ℓ_task_ was defined by the discrepancy between the model output and the rule-generated target response. For neural-behavioral decoding datasets, ℓ_task_ was defined according to the prediction target, including behavioral choice, licking event or stimulus category. In real mouse neural behavioral decoding experiments, motif-frequency constraints including Ave, MOp and FRP were compared with a structure-free Mamba baseline.

### Baseline models

We compared DigiMus with several sequence-modeling baselines commonly used for temporal neural and behavioral data. The baseline models included a temporal convolutional network (TCN), a long short-term memory network (LSTM), a Transformer encoder and a structure-free Mamba model. These baselines were selected to cover convolutional, recurrent, attention-based and state-space sequence modeling strategies.

The TCN baseline used one-dimensional causal convolutions with dilated convolutional layers to expand the temporal receptive field while preserving temporal ordering. The LSTM baseline used gated recurrent units with input, forget and output gates to model temporal dependencies and reduce vanishing-gradient effects in long sequential inputs. The Transformer baseline used self-attention layers to model pairwise dependencies across time steps. The structure-free Mamba baseline used the same temporal state-space backbone as the DigiMus implementation, but did not include region-specific motif regularization or multi-region-capable structural organization.

For all baselines, the input format, output targets, training data, validation data and test splits were kept identical to those used for DigiMus. Model width, depth and hidden dimensions were adjusted to keep the overall parameter scale comparable across models whenever possible. All baseline models were trained using the same task loss as DigiMus, but without connectome-derived motif regularization, region-specific CIS blocks or multi-region-capable organization. This design allowed us to evaluate whether the performance differences arose from the connectome-informed spiking and motif-regularized components rather than from differences in data preprocessing or task definition.

### Cognitive-task evaluation and statistical analysis

DigiMus and baseline models were evaluated on rule-based cognitive tasks using the same input-output task format, train-validation-test protocol and performance metric. The cognitive-task benchmark contained 18 tasks covering Go/Anti sensorimotor mapping tasks and decision-making tasks. In **Fig. 3**, six representative tasks were shown for compact visualization: React Go, FD Go, DMS Go, DM1, Multi DM and Multi delay DM.

For each task, DigiMus was compared with three sequence-modeling baselines: temporal convolutional network (TCN), long short-term memory network (LSTM) and Transformer encoder. The DigiMus bar in **Fig. 3d** corresponds to MotifMamba, the motif-regularized Mamba implementation evaluated under the DigiMus framework. Each model was evaluated across five independent training runs using random seeds 0-4. Task performance was quantified as accuracy on the held-out test set and reported as percentage accuracy.

Bars in **Fig. 3d** show the mean accuracy across five random seeds. Error bars indicate the standard deviation (SD) across the same five seed-level accuracies. Statistical comparisons were performed as exploratory model-level analyses between each baseline model and DigiMus separately for each task using two-sided Welch’s t-tests on the five seed-level accuracy values. These comparisons were used as exploratory pairwise model-level comparisons. Significance levels were annotated as \**P* < 0.05, \*\**P* < 0.01 and \*\*\**P* < 0.001; non-significant comparisons were not annotated. Reported *P* values were nominal and unadjusted, and no multiple-comparison correction was applied for **Fig. 3d**, these P values should therefore be interpreted descriptively rather than as confirmatory statistical evidence.

### Internal dynamics analysis

To quantify the internal dynamics shown in **Fig. 4**, we extracted the hidden state and output state of the trained DigiMus model from held-out test trials for each task included in the **Fig. 4** internal-dynamics analysis. For task *k*, h_*k,i,t*_ denotes the hidden-state vector and *y*_*k,i,t*_ denotes the output-state vector of trial *i* at Mamba step *t*. The number of test trials is denoted by *N*_*k*_, and the hidden-state and output-state dimensions are denoted by *D*_h_ and *D*_*y*_, respectively.

For each task and each time step, the cross-trial mean hidden and output states were computed as

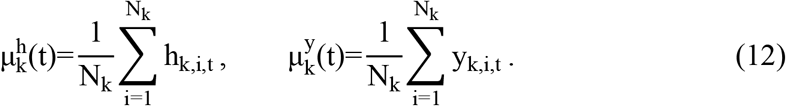

The *h-dev* and *y-dev* trajectories in **Fig. 4a** were defined as signed population-averaged deviations from the corresponding time-specific cross-trial mean. We denote them as Δh_*k,i*_(*t*) and Δ*y*_*k,i*_(*t*):

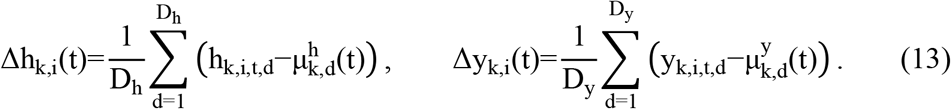

Because the deviations are centered by the time-specific cross-trial mean, their cross-trial average at each time step is zero up to numerical precision. Trial-to-trial bandwidth was therefore quantified at each time step as

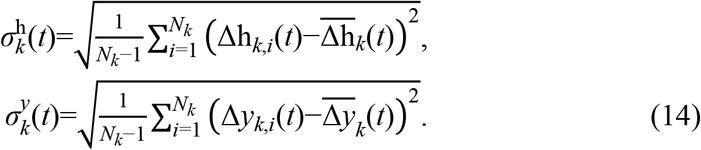

Where,

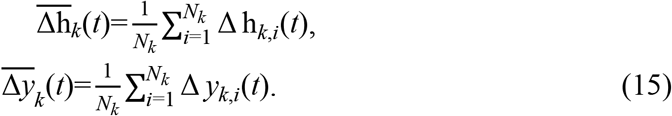

For visualization in **Fig. 4a**, individual trial trajectories were plotted together with dispersion envelopes defined as

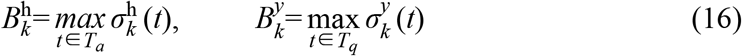

Task error rate was defined as

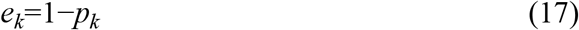

Where *p*_*k*_ denotes the task performance shown above each **Fig. 4a** panel. Associations between task error rate and peak bandwidth were evaluated across the six representative tasks, with each task treated as one effective sample (*n*=6). The five random seeds were not treated as independent samples for this task-level association analysis.

Linear trends in **Fig. 4b,c** were fitted separately for hidden-state and output-state bandwidth:

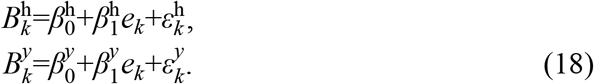

Pearson correlation, R^2^, Spearman rank correlation and corresponding P values were reported as exploratory association statistics. Bonferroni correction was applied across the *h-dev* and *y-dev* association tests.

### Visualization software for DigiMus inspection

The DigiMus inspection interface was implemented as a modular visualization layer for linking model outputs with anatomical and behavioral displays. The software supports brain-atlas exploration, motif-prior configuration, region-specific CIS block assignment, activity visualization and behavioral-output inspection. Registered mouse-brain structures, projection references and motif-related priors can be loaded as structural references, while model-generated activity summaries can be displayed as temporal traces, spatial maps or sectional views.

The simulation-facing component was linked to MuJoCo-based modules, with three-dimensional anatomical and behavioral assets prepared using Blender and SolidWorks according to the visualization and experimental requirements. A communication interface was used to connect the DigiMus controller with the simulation environment, receive model-generated action commands and return behavior-related outputs for inspection. This software layer was used to support model visualization, structure-function inspection and simulation monitoring, rather than to provide independent validation that DigiMus fully reproduces the biological mouse brain.

## Acknowledgments

The authors thank the Brain Science Data and Computing Center and the Digital Brain platform for providing data resources and computational support. The authors also thank members of the laboratory for discussions on digital brain modelling, neural-behavioural decoding and visualization-system design. **Funding** This work was supported by the Brain Science and Brain-like Intelligence Technology - National Science and Technology Major Project (2025ZD0217200), Strategic Priority Research Program of the Chinese Academy of Sciences (Grant No. XDB1010302), CAS Project for Young Scientists in Basic Research (YSBR-116), Youth Innovation Promotion Association CAS, Shanghai Leading Talent Program of Eastern Talent Plan, the Lingang Laboratory Fund (Grant No. LG-GG-202402-06-07, LGL-1987-09), the Shanghai Municipal Science and Technology Project (Grant No. 25ZR1401370, 25LN3200400), Special Support Project of Guangdong Province (Grant No.0720240209). The numerical calculations in this study were carried out on the ORISE Supercomputer. **Author contributions**: Y.L. drafted the manuscript, study implementation, prepared the figures, organized the results and coordinated manuscript revision. X.Z. provided data support and visualization support. Y.S. contributed to model design, algorithm development and methodological discussion.

W.Y. and J.Z. contributed to experimental support, result generation and result verification. X.C. and C.H. contributed to figure preparation, visualization refinement and result organization. T.Z. conceived and designed the study, supervised the project, provided methodological guidance, contributed to scientific discussion and revised the manuscript. All authors discussed the results and approved the final manuscript. **Competing interests:** The authors declare that they have no competing interests.

## Data availability

Public mouse brain resources used in this study are available from CEBSIT, Chinese Academy of Sciences platforms, including the Brain Science Data and Computing Center and the Digital Brain platform (https://digital-brain.cn/). The Allen Brain Observatory Visual Coding Neuropixels dataset is available from the Allen Brain Map data portal. The A1 calcium-imaging and M2 spiking datasets are cited in References [48] and [49]. Processed feature matrices, trained model checkpoints and analysis scripts will be released with a later version of this work or made available from the corresponding author upon reasonable request, subject to institutional data-use policies.

## Code availability

The code used for model construction, motif-frequency regularization, baseline training, neural-behavioural decoding and figure-level analysis will be made available in https://github.com/Ap1rate/DigiMus. Until public release, code and processed analysis scripts are available from the corresponding author upon reasonable request, subject to institutional data-use and software-release policies.

## Supplementary Figure

**Supplementary Fig. 1.**
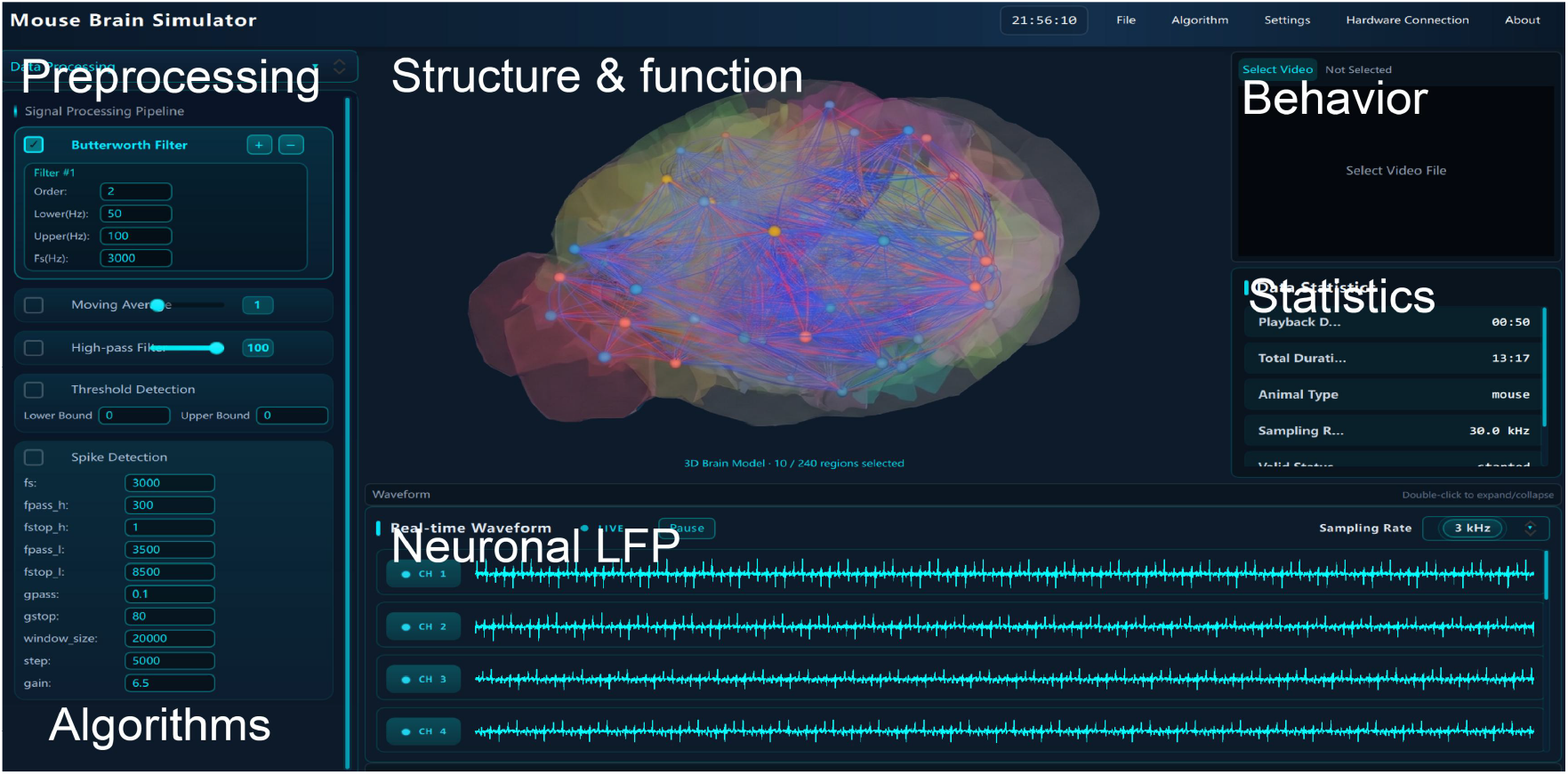
Original DigiMus visualization interface.

**Supplementary Fig. 2.**
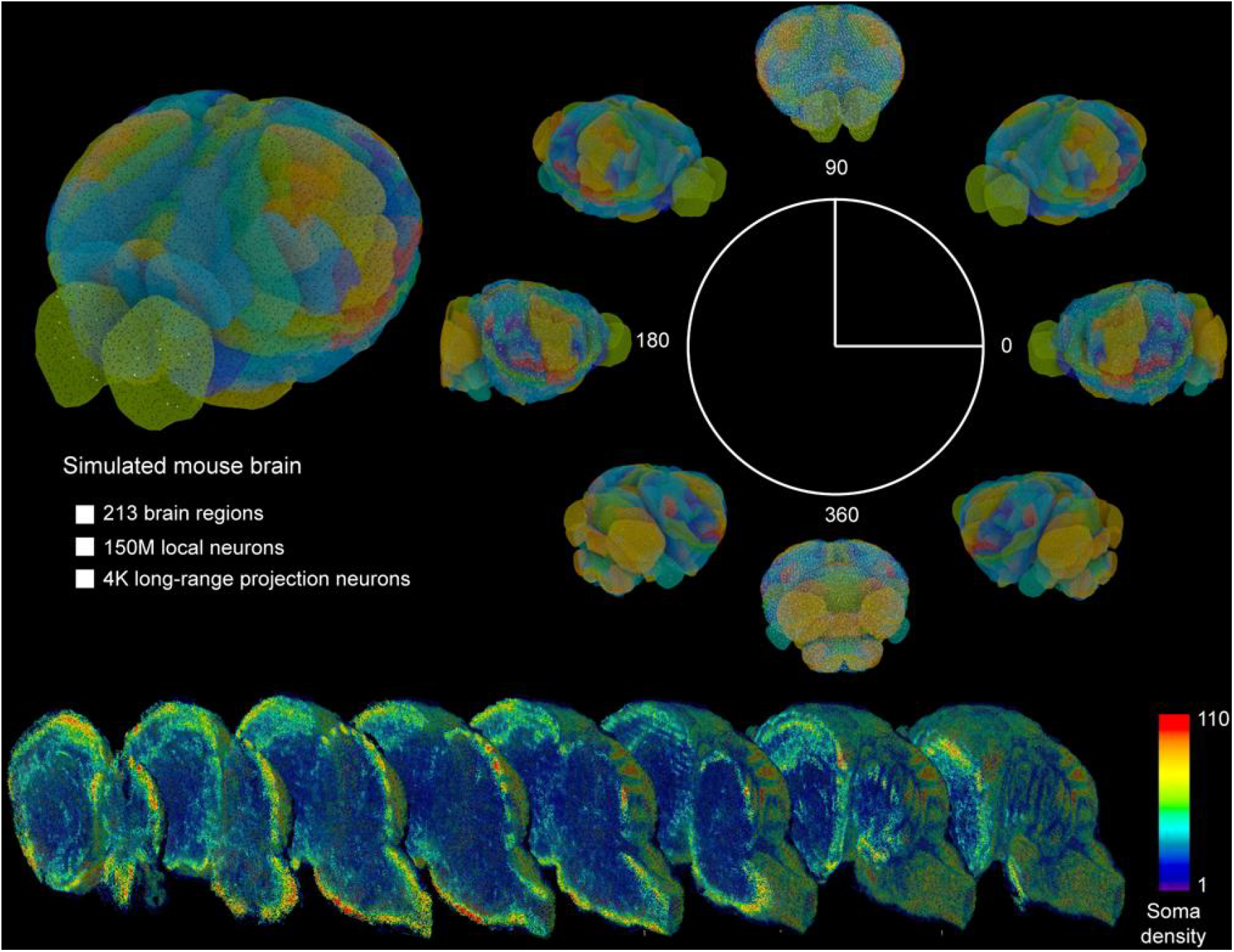
Simulated mouse-brain region and activity visualization.

**Supplementary Fig. 3.**
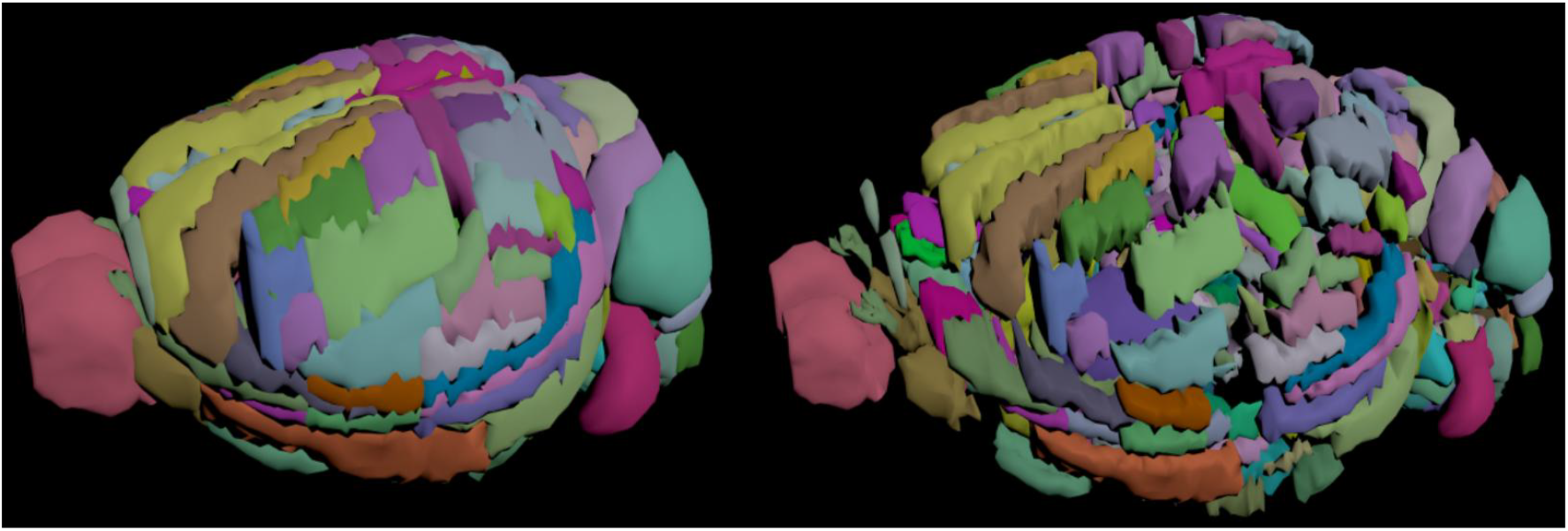
Registered mouse-brain atlas views used for visualization.

## Supplementary Tables

**Supplementary Table 1.** Statistical reporting used for Fig. 3d.

| Item | Reporting choice |
| --- | --- |
| Representative tasks | React Go, FD Go, DMS Go, DM1, Multi DM and Multi delay DM |
| Compared models | TCN, LSTM, Transformer encoder and DigiMus (MotifMamba implementation) |
| Independent runs | Five random seeds per model and task, seeds 0-4 |
| Metric | Test-set accuracy, reported as percentage accuracy |
| Bar height | Mean accuracy across five seeds |
| Error bar | SD across five seeds |
| Statistical test | Two-sided Welch's t-test comparing each baseline with DigiMus |
| Significance threshold | *P < 0.05, **P < 0.01 and ***P < 0.001 |
| Multiple-comparison correction | Not applied; displayed P values are nominal and unadjusted |

**Supplementary Table 2.** Association between task error rate and peak trial-to-trial bandwidth in Fig. 4b,c.

| Item | y-dev | h-dev |
| --- | --- | --- |
| Effective sample size | n = 6 tasks | n = 6 tasks |
| Bandwidth summary | peak max $\sigma(t)$ | peak max $\sigma(t)$ |
| Slope | $6.7 \pm 4.0$ | $3.0 \times 10^3 \pm 3.1 \times 10^3$ |
| Pearson r | 0.64 | 0.43 |
| R<sup>2</sup> | 0.41 | 0.18 |
| Pearson P | 0.17 | 0.4 |
| Spearman $\rho$ | 0.7 | 0.7 |
| Spearman P | 0.12 | 0.12 |
| Bonferroni-corrected P | $\geq 0.25$ | $\geq 0.25$ |
| Interpretation | positive, non-significant trend | positive, non-significant trend |

**Supplementary Table 3.**
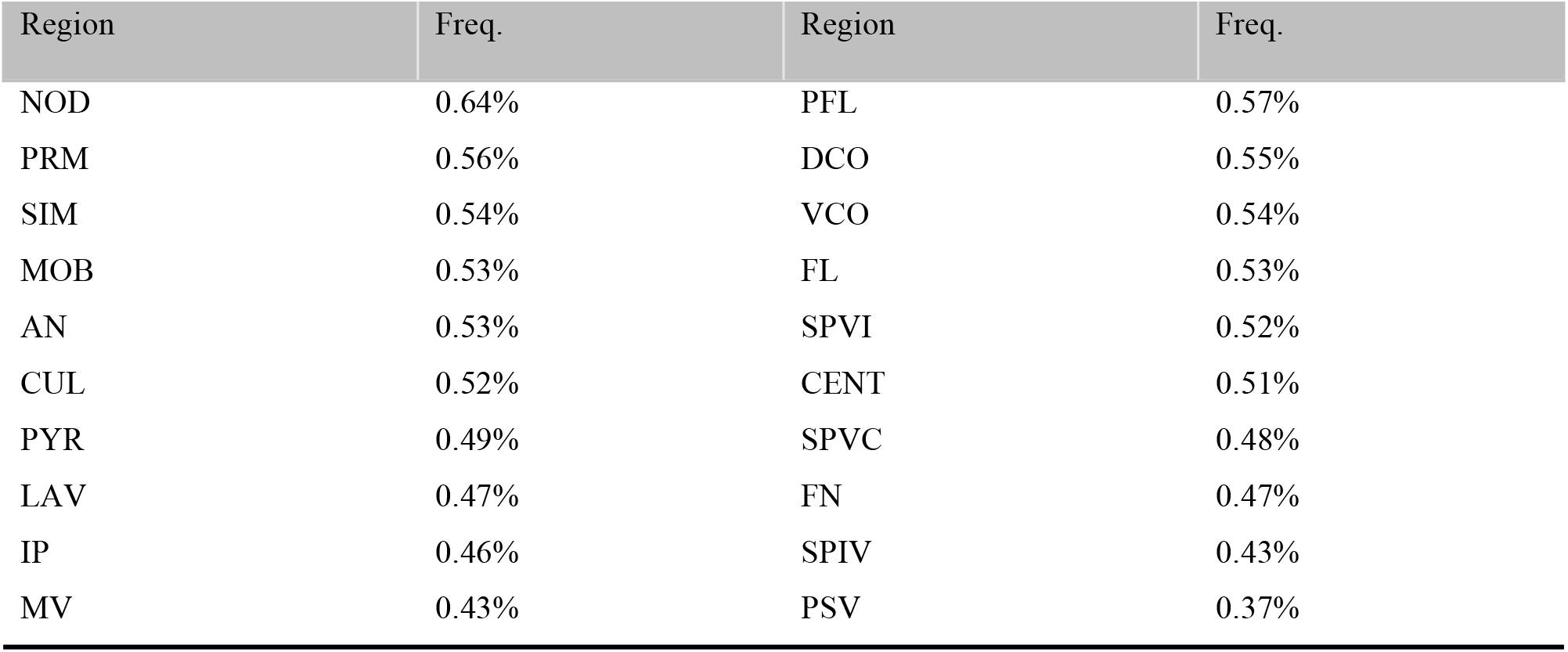
Regional motif-frequency priors used in Fig. 5g-i.

